# Targeting AML Resistance with Two Novel Combinations Demonstrate Superior Efficacy in *TP53, HLA-B, MUC4* and *FLT3* mutations

**DOI:** 10.1101/2024.12.31.630711

**Authors:** Elham Gholizadeh, Ehsan Zangene, Alun Parsons, Mika Kontro, Caroline A. Heckman, Mohieddin Jafari

**Author notes:** Corresponding author: Mohieddin Jafari.

## Abstract

Acute myeloid leukemia (AML) is a genetically heterogeneous malignancy characterized by the clonal expansion of myeloid precursor cells. Despite the advent of venetoclax-based regimens, resistance mechanisms remain a major clinical challenge, particularly in patients with high-risk mutations such as *TP53, MUC4, HLA-B and FLT3*. This study aims to investigate the efficacy of drug combinations for the treatment of AML using both, AML cell lines and zebrafish embryo xenograft model. Specifically, we focus on two drug combinations; the pan-RAF inhibitor LY3009120 combined with the mTOR inhibitor sapanisertib (designated as LS), and the JAK1/2 inhibitor ruxolitinib combined with the ERK inhibitor ulixertinib (designated as RU). The study integrates real-time cell viability assays, xenograft imaging, and genomic analyses to assess drug efficacy and explore correlations between treatment responses and mutational profiles, particularly *TP53, MUC4, HLA-B and FLT3* mutations. Both combinations demonstrated superior efficacy compared to venetoclax-based therapy, with LS notably reducing viability in MOLM-16 and SKM-1 cells, and RU showing comparable efficacy with a favorable safety profile. In zebrafish embryos, LS combination effectively inhibited the proliferation of xenografted human AML cells, as evidenced by decreased fluorescence signals, indicating cell death. The RU combination also disrupted survival of cancer cells, showing promise as a therapeutic strategy. Furthermore, a correlation was identified between drug response and mutational profiles, with *TP53, MUC4, HLA-B and FLT3* mutations significantly influencing sensitivity to the LS and RU combinations. These findings support the further development of LS and RU as effective alternatives to current clinical regimens, with implications for personalized AML treatment.

## Introduction

Acute myeloid leukemia (AML) is a heterogeneous hematological malignancy characterized by the clonal proliferation of myeloid precursor cells, leading to impaired hematopoiesis^1^. The molecular landscape of AML is highly diverse, comprising recurrent somatic mutations that influence disease pathogenesis, treatment response, and clinical outcomes^2^. Advancements in next-generation sequencing (NGS) have facilitated the identification of key driver mutations, enabling risk stratification and the emergence of precision medicine approaches in AML management ^3^.

Mutations in the *TP53* and *FLT3* are among the most critical genetic mutations in AML and play a pivotal role in both prognosis and treatment selection ^4,5^. The *TP53* gene, located on chromosome 17p13.1, encodes the p53 protein, a pivotal tumor suppressor that plays a fundamental role in cellular responses to stress, including DNA damage and oncogenic signaling ^6,7^. Mutations in *TP53*, which occurs in a significant proportion of patients, lead to impaired responses to chemotherapy and poor patient outcomes in AML ^8,9^. The initial TCGA (The Cancer Genome Atlas) report found an 8% prevalence of *TP53* mutations in newly diagnosed AML, with affected patients showing low response rates to standard therapies, high relapse rates, and poor survival outcomes ^10^. Interestingly, *TP53* mutations are often the sole genetic alteration in up to 75% of AML cases, with a reduced incidence of other mutations such as *FLT3, NPM1, IDH, DNMT3A* and *RAS*, suggesting a distinct mutational profile ^10–13^. Induction chemotherapy remains standard for fit AML patients but is less effective in *TP53*-mutated patients, with low response rates (20–40%), high relapse rates, and a median survival of 4–9 months ^14,15^. However, untreated patients have even worse outcomes, highlighting the aggressive nature of *TP53*-mutated AML and the necessity of treatment ^5,16^.

*FLT3* is another gene frequently mutated in AML, making it a critical target for therapeutic intervention. Mutations in *FLT3*, particularly internal tandem duplications (ITD), are associated with poor prognosis and decreased overall survival in AML patients ^17^. The overexpression of *FLT3* leads to increased proliferation of leukemic cells, contributing to the disease’s aggressive nature. *FLT3* inhibitors have been developed to target these mutations, but resistance often develops, necessitating biomarkers to stratify patients and guide treatment decisions ^18^.

Emerging evidence has also highlighted the roles of *MUC4* and *HLA-B* in AML. Overexpression of *MUC4*, a membrane-bound mucin protein, has been linked to chemoresistance and inferior survival outcomes ^19^. Recent findings indicate that *MUC4* expression is significantly higher in AML patients than in healthy controls, correlating with shorter disease-free intervals and reduced overall survival ^20^. This antigen presentation is essential for the recognition and elimination of infected or malignant cells ^21^. Alongside this, HLA-B is a critical component of the human leukocyte antigen (HLA) system, which plays a pivotal role in the immune response by presenting peptides to cytotoxic T cells ^22^. This antigen presentation is essential for the recognition and elimination of infected or malignant cells ^21^. The HLA-B*44 allele, which carries a Bw4-80T motif, weakly binds to the inhibitory NK receptor KIR3DL1, potentially impairing NK cell responses. This weak interaction has been associated with diminished immunotherapy efficacy and lower survival rates in AML, highlighting the importance of HLA-B–NK receptor interactions in cancer treatment ^23^.

Given the high prevalence of genetic mutations and the associated treatment resistance in AML, personalized therapies based on mutational profiles are critical ^24^. While venetoclax, a BCL-2 inhibitor, has shown promise in combination with hypomethylating agents or low-dose cytarabine, resistance, particularly in TP53-mutated or high-risk subtypes, continues to limit its success ^25^. Additionally, discrepancies between *in vitro* and *in vivo* outcomes present ongoing challenges in evaluating drug efficacy. To address this, zebrafish have emerged as a valuable *in vivo* model organism for cancer research, particularly in the context of xenograft studies. The transparent nature of zebrafish embryos allows for real-time imaging of cancer cell behavior, providing insights into tumor growth, invasion, and response to therapy ^26,27^. Furthermore, the compatibility of zebrafish with high-throughput drug screening enhances its utility in evaluating the efficacy of therapeutic combinations. By transplanting human AML cell lines into zebrafish embryos, researchers can investigate the *in vivo* effects of drug treatments, thereby bridging the gap between in vitro studies and clinical applications ^28,29^.

To address the unmet need for effective therapies in high-risk AML,, we investigated two recently proposed drug combinations, developed by our research group^30^, in both AML cell lines and xenografted zebrafish embryos. The drug combinations are pan-RAF inhibitor LY3009120 in combination with the mTOR inhibitor sapanisertib (designated as LS), and the JAK1/2 inhibitor ruxolitinib in combination with the ERK inhibitor ulixertinib (designated as RU). These combinations target key signaling pathways involved in AML progression, the RAF-MEK-ERK pathway which is known to be hyperactivated in AML, and plays a crucial role in cell proliferation and survival ^31^. By targeting this pathway with a pan-RAF inhibitor, we aim to disrupt the signaling cascade that promotes leukemic cell growth. Additionally, the mTOR pathway is integral to cell metabolism and growth; inhibiting this pathway in combination with RAF inhibition may yield synergistic effects, particularly in venetoclax-resistant AML subtypes. Furthermore, the JAK/STAT signaling pathway has been implicated in the pathogenesis of AML, with ruxolitinib demonstrating efficacy in reducing leukemic cell proliferation ^32^. The addition of ulixertinib, an ERK inhibitor, is expected to enhance the therapeutic effect by simultaneously targeting multiple signaling pathways involved in AML progression.

By evaluating the effects of these combinations on a panel of AML cell lines and xenografted zebrafish embryos, we assessed their potential as alternatives to current treatments, particularly in cases with mutations in *TP53, FLT3, HLA-B* and *MUC4*. Our results demonstrate that both LS and RU combinations not only offer superior efficacy compared to venetoclax-based therapies but also show promising safety profiles, indicating their potential for further clinical exploration.

## Results

### Comparative Efficacy of Combination Therapies

To comprehensively assess the therapeutic potential of our proposed drug combinations, we designed a multi-step experimental workflow, providing a multidimensional perspective on anticancer activity and optimization strategies (**Fig. 1**). The initial phase of the study employed the RealTime-Glo (RTG) cell viability assay across a panel of ten AML cell lines—MOLM-16, SKM-1, MV4-11, MOLM-13, NOMO-1, KG-1, Kasumi-1, HL-60, PL-21, and ML-2—over a 72-hour time course, with viability measurements taken at 12-hour intervals. Our primary focus was to evaluate two novel therapeutic combinations: LY3009120 (a pan-RAF inhibitor) with Sapanisertib (an mTOR inhibitor), referred to as LS, and Ruxolitinib (a JAK1/2 inhibitor) with Ulixertinib (an ERK inhibitor), referred to as RU^30^. Given that venetoclax-based regimens have become a cornerstone in the treatment of AML, particularly due to their efficacy in patients who are older or unfit for intensive chemotherapy ^33^. Venetoclax has demonstrated significant clinical benefits, especially when used in combination with hypomethylating agents or low-dose cytarabine ^34^. However, despite its effectiveness, resistance to venetoclax-based therapies is a major issue, limiting their long-term success. This resistance is often observed in patients with poor prognostic mutations such as TP53 and FLT3 ^35^. By comparing new drug combinations with venetoclax, we aim to explore potentially more effective alternatives that could offer better efficacy in overcoming the limitations associated with venetoclax-based treatments.

**Figure 1.**
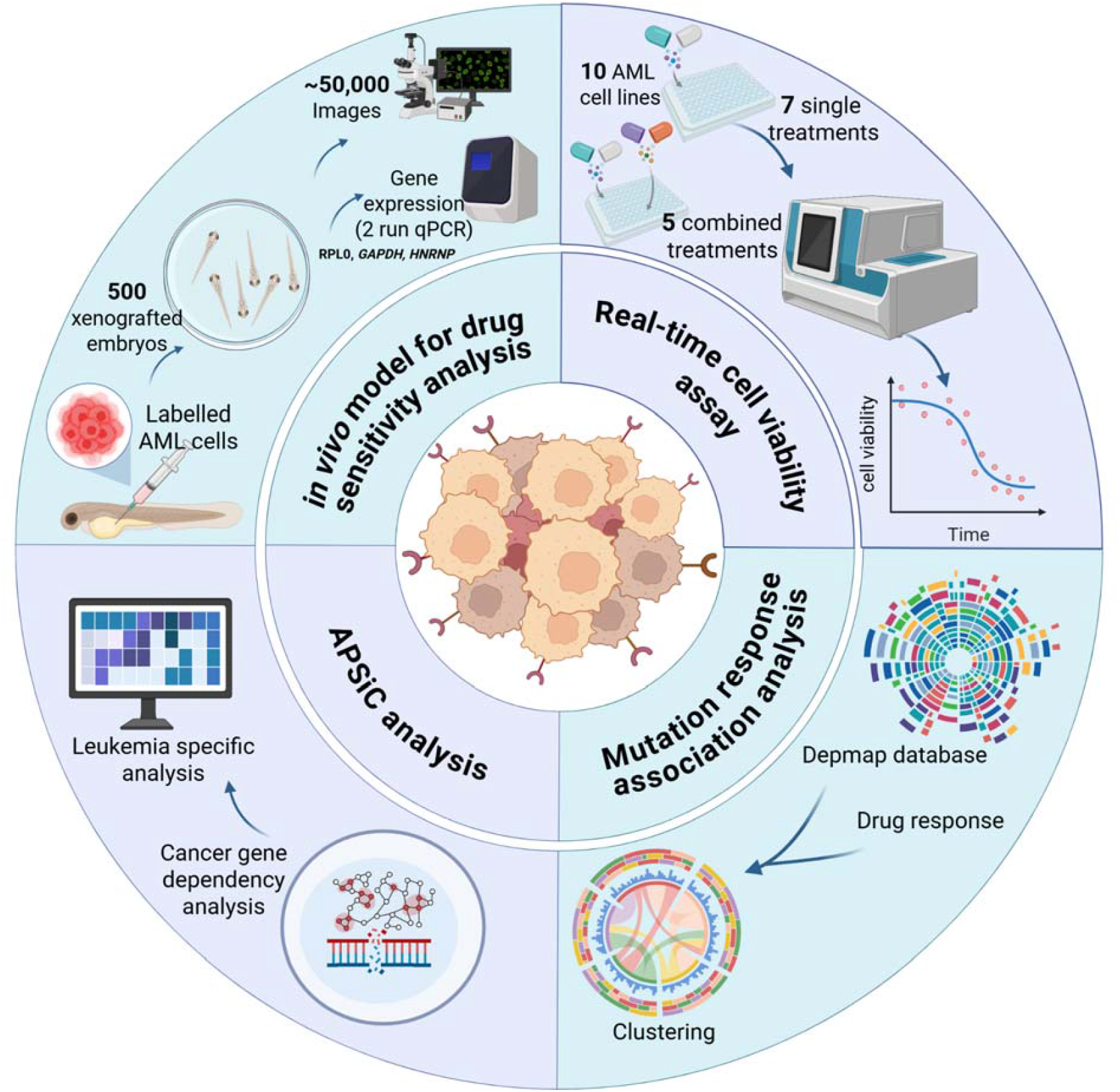
Schematic overview of experimental approaches used to evaluate anticancer treatments across multiple platforms. **Real-time cell viability assay:** Evaluation of 10 AML cell lines over 72 hours, comparing the effects of seven single-agent treatments and five combination regimens. **Mutation response association analysis:** Investigating the correlation between drug sensitivity and specific mutations in various cell lines, with a heatmap to visualize the variation in responses. **APSiC analysis:** Integration of multiomics data to assess gene dependency and mutation relevance across both pan-cancer and leukemia-specific contexts. **In vivo model for drug sensitivity analysis:** Utilizing 500 zebrafish embryos to study the impact of drugs on labeled human cancer cells, with subsequent gene expression analysis and imaging post-treatment.

Detailed cell line-specific analysis revealed that subsets of AML cells were particularly sensitive to the LS combination, achieving significantly greater reductions in cell viability (**Fig. 2A** and **B**, **supplementary Figure S1**). As expected, the combination treatments demonstrated superior efficacy compared to single-agent treatments, with reductions in cell viability observed after 72 hours **(Supplementary figure S1)**. In **Fig. 2A**, MOLM-16 and SKM-1 serve as representative examples, illustrating the pronounced effect of combination therapies compared to individual agents. To benchmark our novel combinations, we compared their efficacy to venetoclax-based therapies across all ten AML cell lines. Using the average responses from all ten AML cell lines, we observed that both LS and RU combinations demonstrated superior or comparable efficacy relative to venetoclax-based therapies (VC, VA, and VR) (**Fig. 2B**). Tukey’s post-hoc analysis showed that the RU regimen performed comparably to the VR and VC regimens (p > 0.6), and demonstrated significantly greater efficacy than the VA regimen (p = 0.014) (**Fig. 2B**). The efficacy of the LS combination is particularly pronounced, with a steep decline in cell viability observed consistently over the 72-hour observation period. This decline suggests a robust anti-leukemic effect that may be attributed to the simultaneous targeting of the RAF and mTOR signaling pathways, which are known to play critical roles in AML cell proliferation and survival. Similarly, the ruxolitinib/ulixertinib combination, targeting the JAK/STAT and ERK pathways respectively, shows comparable efficacy to venetoclax-based combinations.

**Figure 2.** Efficacy and statistical analysis of LS and RU combinations compared to venetoclax-based regimens in AML cell lines. **(A)** Comparative analysis of LS and RU combinations versus single agents and venetoclax-based regimens in SKM-1 and MOLM-16 AML cell lines. Cell viability (%) was measured over 72 hours. Error bars represent the standard deviations. **(B)** Summary of drug response across ten AML cell lines (MOLM-16, SKM-1, MV4-11, MOLM-13, NOMO-1, KG-1, Kasumi-1, HL-60, PL-21, and ML-2) treated with LS, RU, and venetoclax-based combinations. Statistical comparisons between LS/RU and venetoclax-based regimens were conducted using Tukey’s post-hoc test. **Abbreviations:** LY3009120 (L), Sapanisertib (S), Ruxolitinib (R), Ulixertinib (U), Venetoclax (V), Azacitidine (A), Cytarabine (C), LS = LY3009120 + Sapanisertib, RU = Ruxolitinib + Ulixertinib, VA = Venetoclax + Azacitidine, VC = Venetoclax + Cytarabine, VR = Venetoclax + Ruxolitinib.

These findings indicate that targeting both RAF and mTOR pathways in AML provides synergistic effects, particularly in venetoclax-resistant subtypes. The response to the RU combination, although more uniform across all tested cell lines, also indicated potential for personalized treatment strategies. When compared to single-drug therapies, both combinations yielded superior results in reducing AML cell viability, further validating their potential efficacy in clinical settings.

### Drug Sensitivities Across AML Mutations

To further explore differential drug responses among AML cell lines, we conducted a detailed analysis of cell viability for each cell line individually, comparing responses to the novel combinations (LS and RU) against venetoclax-based regimens. Cell viability wa measured at multiple time points (12 to 72 hours) across different cell lines. The results revealed distinct response patterns in different cell lines, with the LS combination consistently inducing greater reductions in viability across most cell lines. The RU combination also demonstrated strong activity in specific cell lines, particularly in comparison to first-line regimens (**Fig. 3**). Notably, both LS and RU combinations significantly reduced cell viability in NOMO-1 and MOLM-16 cell lines in comparison to all other venetoclax based combinations (**Fig. 3**). In contrast, cell lines such as HL60, Kasumi-1, and ML-2, MV-4-11 exhibited relatively lower sensitivity to RU, highlighting the heterogeneity of drug responses across AML models.

**Figure 3.**
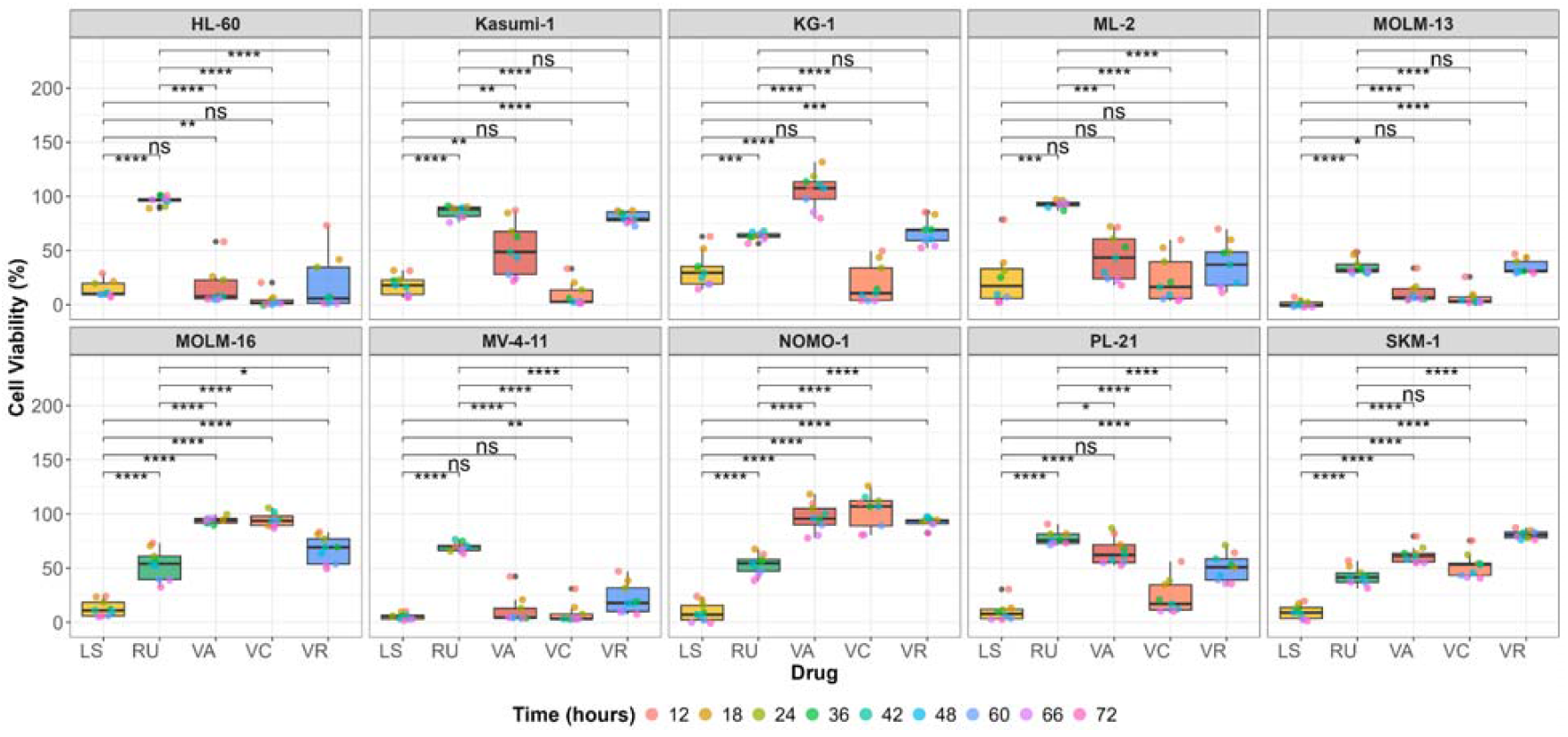
Comparison of cell viability (%) across drug combinations and AML cell lines over time. This plot illustrates the cell viability of multiple AML cell lines treated with five drug combinations: LY3009120 + Sapanisertib (LS), Ruxolitinib + Ulixertinib (RU), Venetoclax + Azacitidine (VA), Venetoclax + Cytarabine (VC), and Venetoclax + Ruxolitinib (VR). Cell viability was measured at multiple time points up to 72 hours. The y-axis represents cell viability (%), while the x-axis denotes the different drug combinations. Boxplots display the distribution of cell viability values for each treatment, with individual data points corresponding to distinct time points. Pairwise comparisons between drug combinations were conducted using two-sample t-tests. Statistical significance is indicated as follows: ***p < 0.001, ****p < 0.0001, and ns (not significant, p > 0.05).

To understand these differential responses in the context of genetic alterations, we performed hierarchical clustering of AML cell lines based on drug combination responses (**Fig. 4**). Upon filtering to include genes mutated in at least three cell lines, MOLM-16 and NOMO-1 emerged as distinct clusters associated with poor responses to venetoclax-based combinations but notable sensitivity to LS and RU.. Both cell lines harbor mutations in *TP53, HLA-B* and *MUC4* which are known to influence treatment responses in AML^36^. *TP53*, a tumor suppressor known as the “guardian of the genome,” is frequently associated with resistance to venetoclax-based therapies ^14,15^.^37^. In our companion work, we have identified *TP53* as a direct target for samples treated with both singles and a combination of ruxolitinib/ulixertinib in lysate cells using the CoPISA method, emphasizing its role in mediating therapeutic response ^38^. *FLT3* mutations are among the most common alterations in AML and have been consistently shown across most studies to be associated with a more aggressive phenotype ^39,40^. Also, patients harbouring this mutation have worse survival and most studies have shown no—or only minimal—prognostic impact of FLT3 mutations ^41,42^. This mutation leads to constitutive activation of FLT3 signaling, promoting uncontrolled proliferation and survival of leukemic cells ^43^. In our analysis, three cell lines, PL-21, MOLM-13, MV-4-11, harbor FLT3 mutations and are mostly sensitive to the combination of VC and LS, with MOLM-13 responding particularly well to LS.

**Figure 4.**
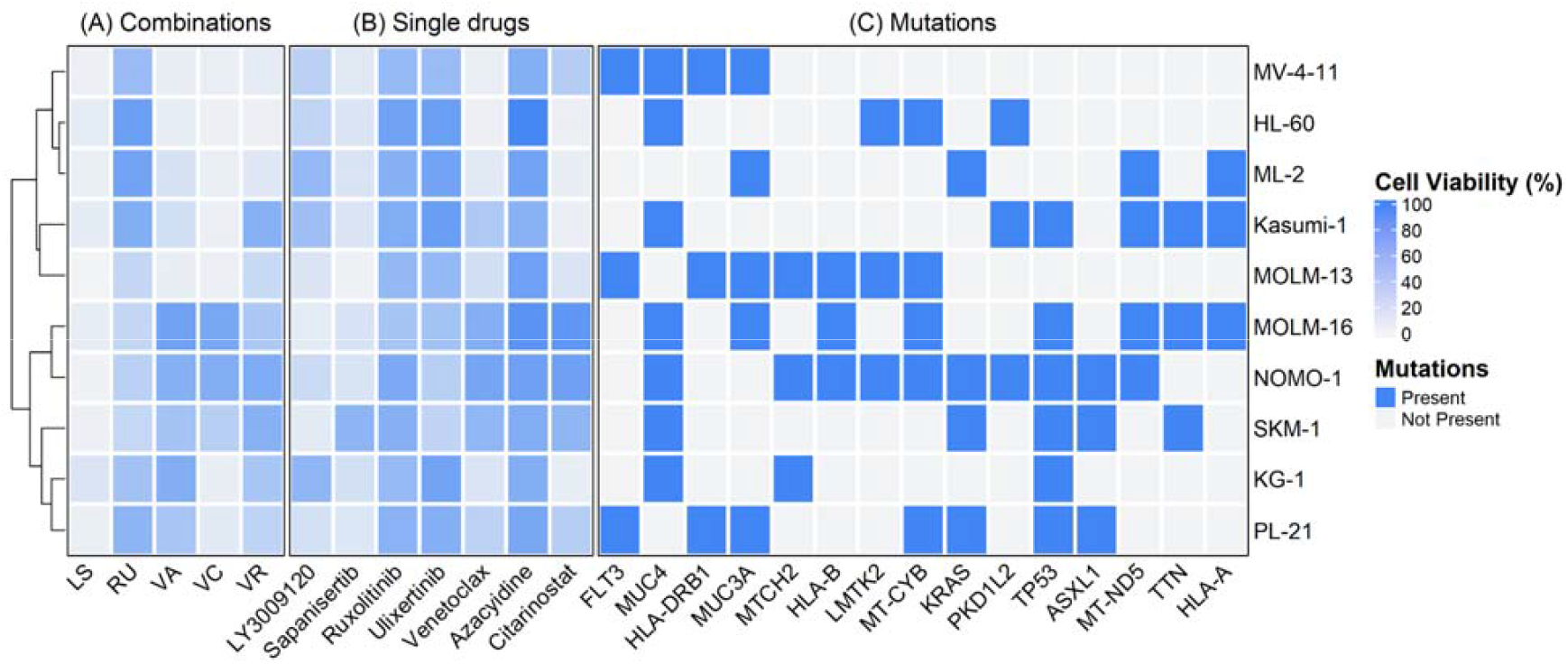
Heatmap of cell viability (%) across drug treatments and mutational profiles. The heatmap shows cell viability after 72-hour treatment of 10 AML cell lines under the following conditions: **(A)** Drug combinations—LY3009120 + Sapanisertib (LS), Ruxolitinib + Ulixertinib (RU), Venetoclax + Azacitidine (VA), Venetoclax + Cytarabine (VC), and Venetoclax + Ruxolitinib (VR); **(B)** Single agents; and **(C)** Mutation profiles, filtered to include only genes mutated in ≥3 cell lines. Hierarchical clustering of rows (cell lines) was performed based on drug combination responses to identify similarities in drug sensitivity patterns. Columns are divided into two main sections: drug combinations and gene mutations. Cell viability is represented by blue color gradient, with darker shades indicating higher viability. Mutation presence is indicated by blue marks at the gene–cell line intersections.

Moreover, we identified *MUC4* mutations in the context of our study, which may influence the response to leukemic cells, further complicating treatment efficacy. *MUC4*, a mucin protein, has been extensively studied for its role in various solid malignancies, where it is linked to tumor growth, proliferation, adhesion, invasion, inhibition of apoptosis, and chemoresistance ^19^. However, its expression and clinical significance in AML have not been thoroughly explored until recently. *MUC4* expression levels are significantly elevated in the bone marrow of AML patients compared to healthy controls. This overexpression is associated with worse survival outcomes and shorter disease-free survival, making *MUC4* an independent prognostic marker for AML, also highlighting its importance in understanding disease progression and treatment response ^20^.

The other mutation in MOLM-16 and NOMO-1 is human leukocyte antigen *HLA-B* which influences the immune system’s ability to recognize and respond to leukemic cells, thus playing a role in both the susceptibility and progression of AML ^44^. Importantly in our previous study ^38^, this gene has been targeted by both combinations and single drugs in lysate and living cells. The presence of these mutations underscores the necessity of targeting specific pathways using these drug pairs, which could be more effective in overcoming resistance mechanisms commonly associated with AML treatments.

Together, these findings underscore the potential of LS and RU combinations in treating AML subtypes with co-occurrence of *TP53, HLA-B* and *MUC4* mutations, with superior efficacy compared to venetoclax-based regimens. Additionally, the LS combination demonstrated strong efficacy in MOLM-13, a cell line harboring *FLT3* mutation. Given the clinical significance of *FLT3*, we selected MOLM-13 along with NOMO-1 for further mechanistic investigations.

### Genomic Alterations and Proteogenomic Correlations

Our integrative analysis highlights key genomic alterations in AML cell lines that correlate with differential responses to targeted therapies (**Fig. 5**). The OncoPrint in Fig.5 displays the top 40 most recurrent mutations, revealing frequent alterations in genes such as *TP53, FLT3, HLA-B* and *MUC4. TP53* mutations were observed in 7 out of 10 cell line (SKM-1, KG-1, Kasumi-1, MOLM-16, NOMO-1, PL-21, and others), aligning with its well-documented role in AML pathogenesis ^45^. *MUC4* and *MUC3A*, both mucin family genes, were frequently altered, particularly in HL-60, MV-4-11, PL-21, MOLM-13, and ML-2, with various missense and frameshift variants, suggesting potential involvement in leukemogenesis ^46^. *FLT3* exhibited in-frame insertions in MOLM-13, MV-4-11, and PL-21, known to confer proliferative advantage and poor prognosis in AML ^47^. *HLA* gene mutations were frequently observed, *HLA-B* mutations were detected in four cell lines (MOLM-13, MOLM-16, NOMO-1, PL-21), predominantly as missense variants. Given *HLA-B*’s critical role in antigen presentation, these alterations may contribute to immune evasion in AML ^48^.

**Figure 5.**
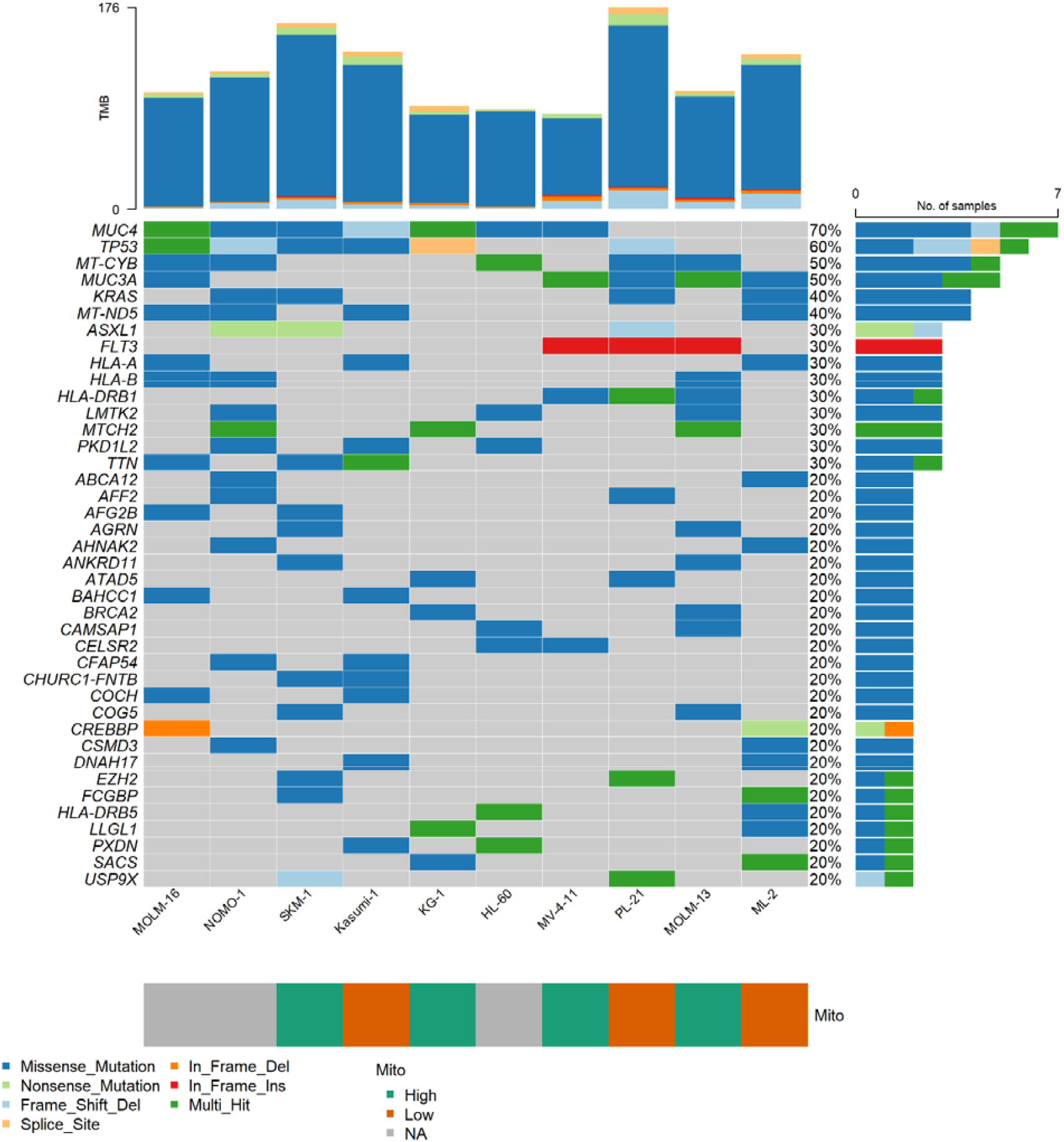
OncoPrint plot showing the most recurrent somatic mutations in 10 AML cell lines. Rows correspond to genomic variants, columns correspond to samples, and variants are ranked by mutation frequency (left). The mutations are filtered to show top 40 mutations. Alterations are color-coded (bottom). The frequency and type of alterations for each sample are shown at the top. Cell lines are classified as Mito low and high based on resources.

The majority of mutations were missense variants (over 70%), indicating amino acid substitutions potentially affecting protein function ^49^. Several frameshift and nonsense mutations were also observed, particularly in tumor suppressors such as *TP53, ASXL1*, and *CREBBP*, likely leading to loss-of-function effects. Mutations in mitochondrial genes, such as *MT-CYB* and *MT-ND5*, were identified in several cell lines including HL-60, ML-2, MOLM-13, and NOMO-1 (**Fig. 5**). These were predominantly missense variants and may reflect alterations in oxidative phosphorylation pathways^50^, relevant in AML metabolic reprogramming ^51^. Recent studies have proposed a stratification of AML into Mito-high and Mito-low subtypes based on mitochondrial protein abundance, which significantly influences therapeutic outcomes ^52^. Mito-high subtypes are characterized by a heightened expression of mitochondrial proteins, which correlates with aggressive disease phenotypes and poor clinical outcomes, particularly a lower remission rate and shorter overall survival post-intensive induction chemotherapy ^52^. This pivotal insight suggests that mitochondrial biogenesis and function play crucial roles in cellular metabolism and survival, substantially influencing treatment responses.

To explore the implications of mitochondrial stratification, we classified AML cell lines into Mito-high (SKM-1, KG-1, MV-4-11, MOLM-13) and Mito-low (ML-2, PL-21, Kasumi-1) groups based on proteomic profiles and analyzed their responses to different drug treatments at the 72-hour time point (**Fig. 6**). As shown in **Fig. 6**, Mito-high cell line consistently demonstrated heightened sensitivity to novel combinations, particularly RU, in comparison to venetoclax-based regimens (VR, VC, VA). In detail, SKM-1, a Mito-high cell line, showed pronounced resistance to venetoclax-based combinations, yet responded robustly to RU and LS with viability reduced to 31% and 1.3%, respectively. This marked shift underscores the potential of RU as a more effective treatment in venetoclax-resistant, Mito-high contexts. Similarly, KG-1 exhibited poor response to VR (52%) and VA (79%) but showed substantial reductions in cell viability under RU (56%) and LS (14%) treatments. Notably, MOLM-13, a FLT3-mutated and Mito-high cell line, showed very low viability under LS treatment. RU also reduced viability to 32%, and also this cell was responding to venetoclax based treatment with VR having highest viability of 29%, further supporting LS’s potency in FLT3-driven and metabolically active AML subtypes.

**Figure 6.**
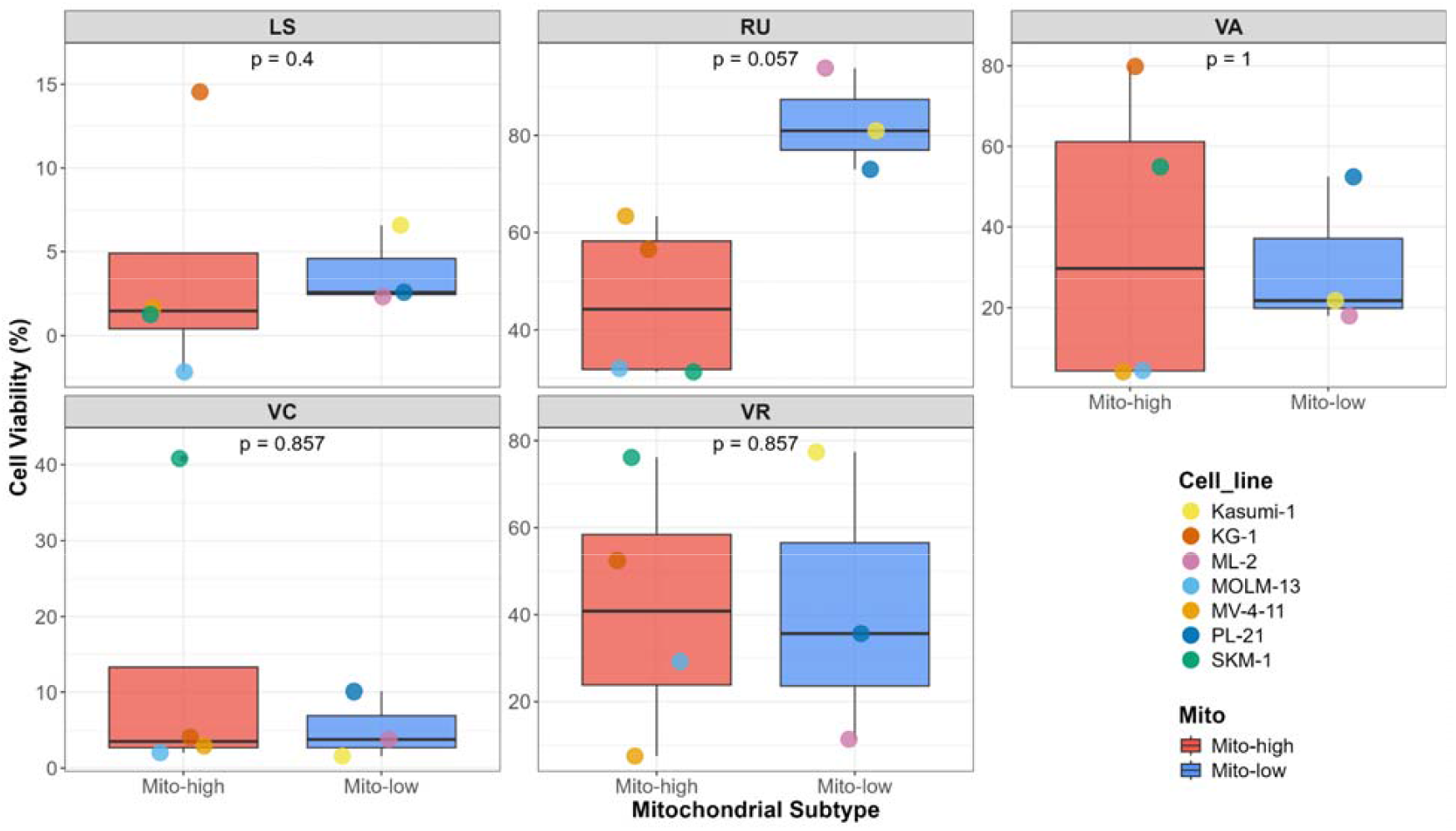
Comparison of cell viability between Mito-high and Mito-low AML cell lines across different drug treatments at 72 hours time point. Boxplots display the distribution of cell viability (%) for each mitochondrial subtype, with individual data points overlaid. Cell lines are categorized as Mito-high (SKM-1, KG-1, MV-4-11, MOLM-13) or Mito-low (ML-2, PL-21, Kasumi-1) based on their mitochondrial profiles. Data points are colored according to the respective cell lines. Statistical significance between Mito-high and Mito-low groups was assessed using the Wilcoxon rank-sum test, and p-values are displayed within each facet.

This integration of genomic and proteomic data offers a more comprehensive understanding of AML heterogeneity. By aligning drug responses with mitochondrial subtypes, we can better predict therapeutic efficacy and tailor combination strategies to overcome resistance, particularly in high-risk Mito-high subtypes. Our findings advocate for the inclusion of mitochondrial profiling in future stratified clinical trial designs, paving the way for more effective and personalized AML therapies.

### APSiC Cancer Gene Dependency Analysis

To further investigate the significance of genomic alterations in the selected AML cell lines, we utilized APSiC, a cancer gene dependency analysis tool. APSiC integrates multiomics and cancer vulnerability datasets, primarily sourced from DepMap, and provides a user-friendly online interface ^53^. Among the hypermutated genes identified in the chosen AML cell lines, *TP53, MUC4*, and *HLA-B* were notably overrepresented. Using APSiC, we evaluated these genes across both pan-cancer and leukemia-specific contexts.

As illustrated in **Fig. 7**, *TP53* was identified as a significantly mutated cancer gene in the pan-cancer analysis, with a p-value approaching zero (p-value = 7.3e-56). However, in the leukemia-specific context, APSiC did not reveal significant missense mutation enrichment (p = 0.37). Conversely, *HLA-B*, a gene with hypermutation potential, exhibited a p-value of 0.054 in the pan-cancer analysis, but the leukemia dataset lacked sufficient samples for statistical validation. The discrepancies observed in APSiC analyses between pan-cancer and AML-specific contexts underscore the necessity of context-dependent experimental studies to identify optimal targeted therapies, whether as single agents or in combination.

**Figure 7.**
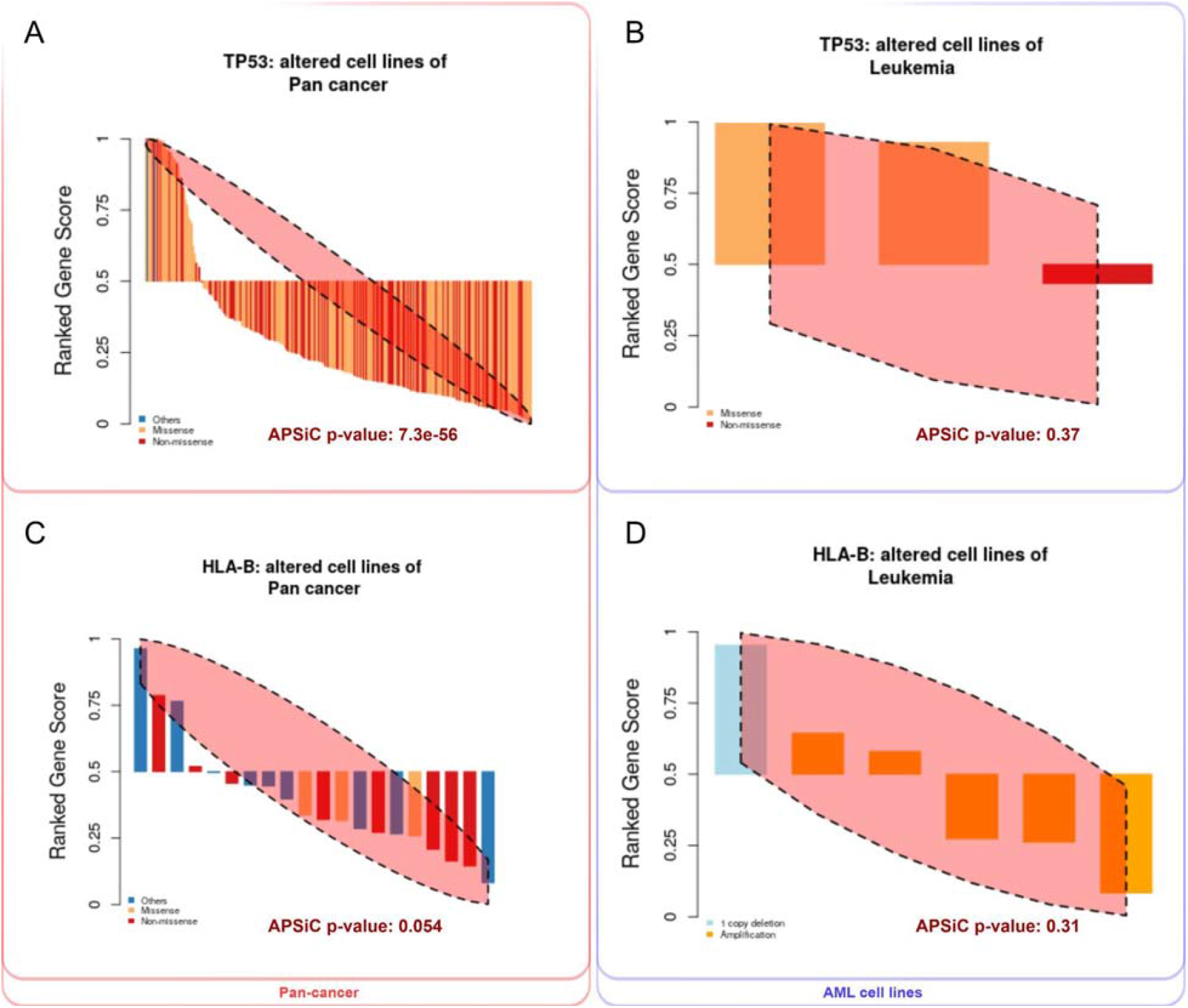
Waterfall plot of CRISPR gene dependency scores (ATARIS scores) for selected cancer cell lines. The y-axis represents cell viability based on CRISPR gene dependency scores, while the x-axis shows ranked cell lines of the cancer types relevant to the study. The dependency scores illustrate the sensitivity of AML cell lines to gene knockouts, highlighting key vulnerabilities in specific genomic contexts. **(A)** TP53 altered cell lines of the pan-cancer context. **(B)** TP53 altered cell lines of the Leukemia context. **(C)** HLA-B altered cell lines of the pan-cancer context. **(D)** HLA-B altered cell lines of the Leukemia context.

These findings emphasize the critical importance of tailoring therapeutic strategies to the specific genetic backgrounds of distinct cancer types, complemented by drug response analyses, particularly in vivo models, to elucidate resistance mechanisms and improve treatment outcomes.

### Image-based Drug Efficacy in Xenografted Zebrafish Embryos

To extend our *in vitro* findings and evaluate the translational potential of novel drug combinations, we employed a xenografted zebrafish embryo model using MOLM-13 and NOMO-1 AML cell lines., MOLM-13 represents a Mito-high AML subtype harboring an *FLT3* mutation and demonstrates broad sensitivity to all tested drug combinations, while NOMO-1, characterized by *TP53, MUC4*, and *HLA-B* mutations, displays selective sensitivity to the novel LS and RU regimens. These specific cell lines were selected due to the clinical relevance of their genetic alterations and their contrasting drug response profiles.

Building on our prior findings in *FLT3*-mutated patient samples treated with LS and RU combinations ^30^, we aimed to assess whether these responses could be recapitulated *in vivo*. Zebrafish embryos were injected with CellTrace Far Red–labeled MOLM-13 or NOMO-1 cells, followed by exposure to drug regimens for 72 hours. Treatment efficacy was evaluated via brightfield and fluorescence microscopy, enabling visualization of both host morphology and human leukemia cell viability. As shown in **Fig. 8**, the microscopy images provided a comprehensive overview of the overall morphology of the zebrafish embryos after treatment with the various drug combinations. The embryos exhibited distinct morphological changes, which were indicative of the effects of the treatments on both the host organism and the xenografted human leukemia cells.

**Figure 8.**
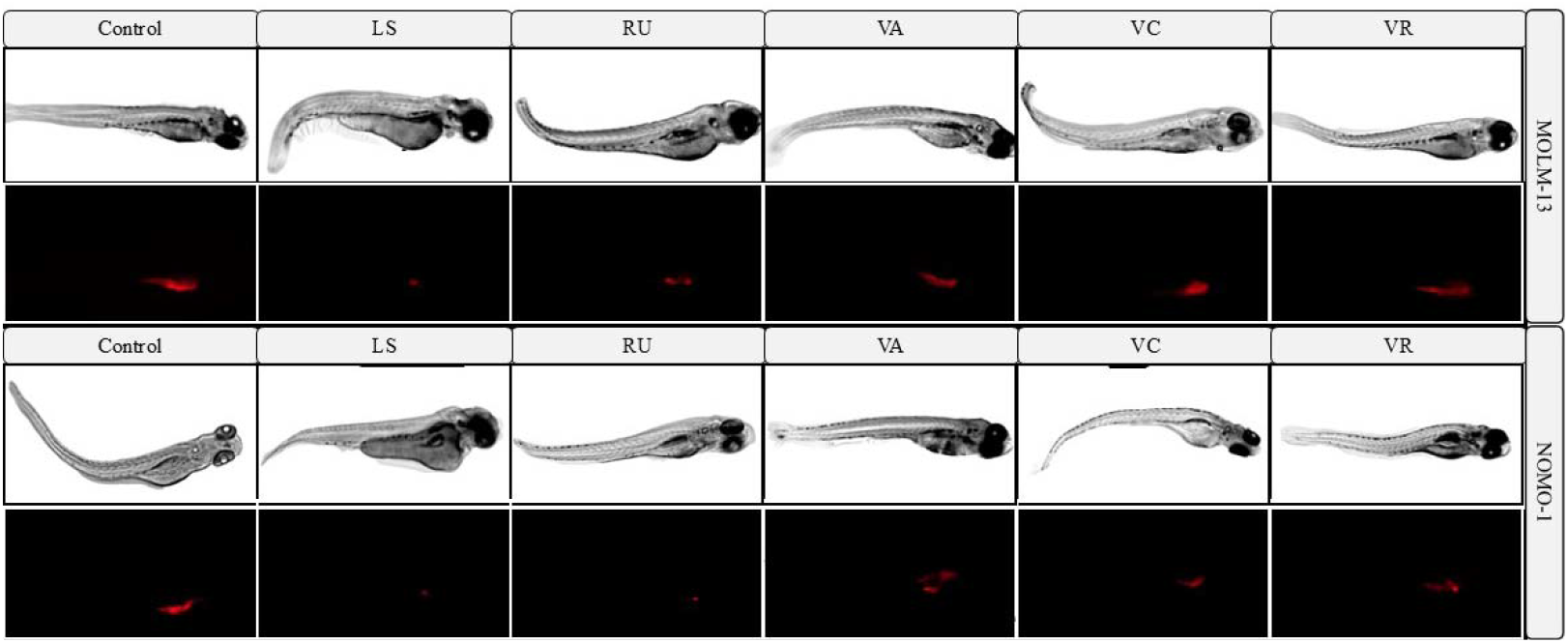
Visualization of drug efficacy in MOLM-13 and NOMO-1 xenografted zebrafish embryos. Brightfield and fluorescence microscopy images of zebrafish embryos injected with MOLM-13 or NOMO-1 cells labeled with CellTrace Far Red, followed by 72-hour incubation and treatment with various drug combinations. Treatments include LY3009120 + sapanisertib (LS), ruxolitinib + ulixertinib (RU), venetoclax + azacitidine (VA), venetoclax + cytarabine (VC), and venetoclax + ruxolitinib (VR). The upper rows display the overall morphology of the zebrafish, while the lower rows highlight fluorescence signals from labeled human cells, reflecting drug effects.

To further quantify these morphological effects and assess potential developmental toxicity, we measured zebrafish embryo lengths using ImageJ software 72 hours post-treatment. Embryo length is a sensitive and reliable marker of developmental progression and systemic stress ^54^. As shown in **Fig. 9A**, xenografted embryos treated with various drug regimens displayed varying degrees of developmental impairment. In the MOLM-13 cohort, untreated controls had an average embryo length of approximately 3 mm. Embryos treated with LS exhibited the most pronounced reduction, followed by VA, RU, and VC, indicating moderate developmental delays. VR treatment resulted in a slightly milder reduction, suggesting a less pronounced impact on host development. NOMO-1 xenografted embryos demonstrated a slightly greater baseline length in controls (mean: 3.1 mm), with drug-treated groups showing more moderate reductions. LS and RU treatments resulted in average lengths of 2.8 mm, while VC and VR treatments maintained embryo lengths around 2.99 mm. The VA combination induced a modest reduction to 2.93 mm. Overall, the developmental impact of drug treatments appeared less severe in NOMO-1–xenografted embryos compared to MOLM-13. Statistical analysis using unpaired t-tests revealed no significant differences in embryo length between treated and control groups for either cell line, suggesting that the observed changes were modest and within a tolerable range. These findings indicate that while the drug combinations exerted therapeutic effects, their impact on host development was limited. Notably, the LS and RU combinations maintained favorable efficacy-to-toxicity profiles, supporting their continued preclinical evaluation.

**Figure 9.** Quantification of leukemic burden and developmental toxicity in zebrafish embryos following combinatorial drug treatment. Human AML cell lines (MOLM-13 and NOMO-1) were labeled with CellTrace Far Red and injected into zebrafish embryos. Following 72 hours of treatment with various dru combinations—LY3009120 + sapanisertib (LS), ruxolitinib + ulixertinib (RU), venetoclax + azacitidine (VA), venetoclax + cytarabine (VC), and venetoclax + ruxolitinib (VR)—embryos were imaged to assess leukemic burden and developmental toxicity. **(A)** Zebrafish embryo lengths were measured using ImageJ software to evaluate potential developmental toxicity. **(B)** Quantification of fluorescence intensity, representing leukemic cell burden, was performed using fluorescence microscopy. Each treatment group was compared to th control group using an unpaired t-test. Data are presented as boxplots for each cell line (MOLM-13 and NOMO-1) and dots represent replicates.

Fluorescence microscopy further confirmed the anti-leukemic efficacy of these regimens. As shown in **Fig. 9B**, signal intensity—reflecting the leukemic burden of labeled human AML cells—was quantified post-treatment. Across both MOLM-13 and NOMO-1 xenografts, combination therapies elicited differential reductions in fluorescence signal compared to untreated controls. In NOMO-1–xenografted embryos, LS and RU treatments resulted in significant reductions in fluorescence intensity, indicating a strong anti-leukemic effect. The VR and VC regimens also reduced signal intensity to 1.40 and 1.30, in contrast, the VA combination showed only a modest and statistically non-significant effect (p = 0.43) relative to the control, suggesting limited activity in this model.

In MOLM-13–xenografted embryos, fluorescence intensity also decreased following combination treatments, although the magnitude of reduction varied. RU significantly lowered leukemic burden, while LS showed a near-significant trend (p = 0.054), reflecting strong cytotoxicity. Other venetoclax-based combinations (VA, VC, VR) resulted in more moderate or variable effects, with non-significant p-values (p = 0.17–0.66) and fluorescence signals ranging from 1.33 to 1.53, relative to the control (1.60).

Collectively, these findings underscore the superior *in vivo* efficacy of the LS and RU combinations in both MOLM-13 and NOMO-1 xenografted zebrafish embryos. These regimens significantly reduced leukemic burden, as evidenced by diminished fluorescence intensity, while maintaining minimal developmental toxicity—highlighting a favorable efficacy-to-toxicity profile. The image-based results are further supported by gene expression analyses, which revealed consistent modulation of key metabolic and structural genes in response to LS and RU. Together, these data strengthen the mechanistic rationale for LS and RU as mutation-informed, clinically translatable therapies for AML. Moreover, venetoclax-based combinations (VA, VC, VR) demonstrated moderate activity, suggesting their utility may depend on specific genetic contexts. Overall, this work supports the continued preclinical development of LS and RU, particularly in genetically heterogeneous AML subtypes.

### Gene expression-based Drug Efficacy in Xenografted Zebrafish Embryos

To assess the molecular effects of novel and first-line drug regimens *in vivo*, we evaluated gene expression changes in MOLM-13 and NOMO-1 xenografted zebrafish embryos. These cell lines harbor key AML-associated mutations—*FLT3* and *TP53*, respectively—and exhibit distinct drug sensitivity profiles. Human leukemia cells were injected into zebrafish embryos, followed by 72-hour drug treatments. Subsequently, total RNA was extracted to enable transcriptomic analysis of the human cancer cells within the xenografted embryos, providing insight into drug-induced gene expression alterations in a physiologically relevant context.

We focused on three human reference genes—RPL0, *GAPDH*, and *HNRNP*— selected for their stability across AML studies ^55–57^. Human gene expression was normalized against zebrafish *RPL13* gene using the ΔΔCt method, allowing accurate quantification of human gene expression in the chimeric embryo context^58^. A two-way ANOVA revealed significant effects of treatment, genes, and their interaction across most sample groups in both cell lines (**Fig. 10** and **Supplementary Table S2**). In MOLM-13 xenografted embryos, treatment was the dominant factor influencing gene expression (e.g., LS: p = 2.26e-10), with notable gene–treatment interactions also detected, highlighting the context-specific modulation of gene expression by therapy. In the NOMO-1 xenografted embryos, the two-way ANOVA results mirrored those observed in MOLM-13, with treatment being a significant factor in gene expression changes **(Supplementary Table S2)**. The LS treatment group showed a significant p-value of 5.48e-10 for treatment effects, indicating its potent influence. However, the gene-treatment interaction was not significant (p = 0.086), suggesting a more uniform effect of LS treatment across the genes studied. The RU group in NOMO-1 exhibited significant treatment effects with a p-value of 1.17e-07. However, the gene-treatment interaction was not significant (p = 0.527), suggesting a consistent treatment effect across different genes. The VA treatment group showed significant effects with a p-value of 3.03e-10, and the gene-treatment interaction was significant (p = 0.000559), indicating a nuanced modulation of gene expression. The VC and VR treatment groups also demonstrated significant treatment effects with p-values of 3.61e-08 and 4.47e-10, respectively.

**Figure 10.**
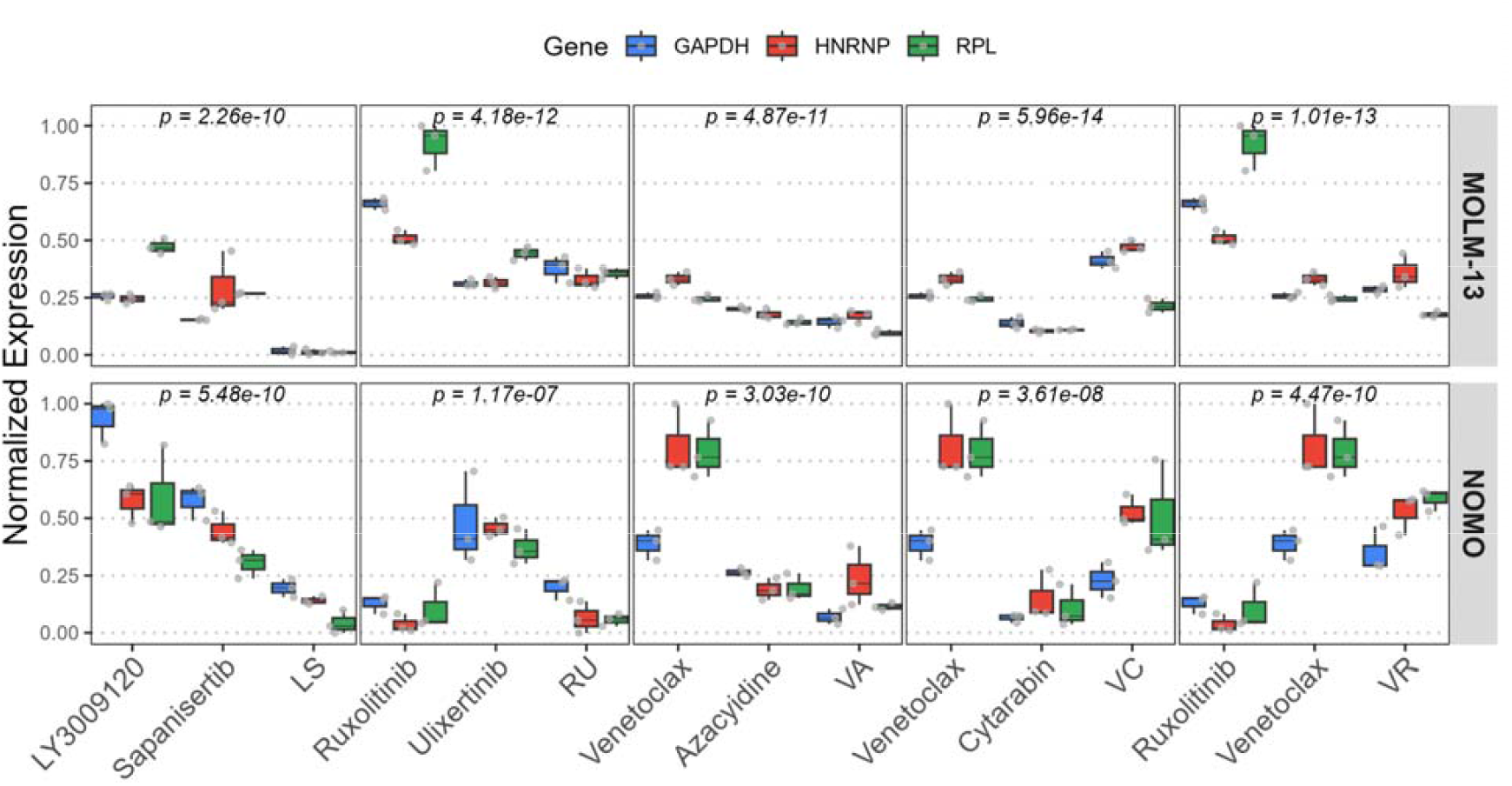
Gene expression of human cells in xenografted zebrafish. This figure illustrates the normalized expression levels of three genes—GAPDH, HNRNP, and RPL— across various drug treatments in MOLM-13 and NOMO-1 cell lines. Human cells were injected into 3 days post-fertilization (dpf) zebrafish embryos, followed by a 72-hour treatment with a panel of single and combination therapies. Each panel corresponds to a treatment condition and shows gene expression variability within and between groups.. Boxplots display log-transformed normalized expression values to the zebrafish reference gene, *RPL13*, using the ΔΔCt method, with individual data points (jittered) to reflect distribution. Genes are color-coded and grouped by treatment and cell line. Statistical analysis was performed using two-way Classical ANOVA, followed by post hoc pairwise comparisons. p-values correspond to the main effect of Treatment and full ANOVA tables available in **Supplementary Table S2** and **S3**.

The pairwise comparison **(Supplementary Table S3)** further supported these findings, indicating robust and gene-specific expression changes in response to drug regimens across both cell lines. For the *GAPDH* gene, significant differences were observed in response to various treatments in both cell lines. In MOLM-13, the LS treatment resulted in a significant decrease in *GAPDH* expression in comparison to LY3009120 treatment (p = 0.000470), whereas RU treatment led to a downregulation in compare to Ruxolitinib (p = 4.29e-07). In contrast, the combinations of VA and VC treatments displayed non-significant changes, highlighting the superior efficacy of LS and RU in modulating *GAPDH* expression. In NOMO-1, the LS treatment also significantly decreased *GAPDH* expression in comparison to single treatments (p = 5.51e-27), while RU treatment showed significant effect in comparison to ulixertinib but non-significant effects in ruxolitinib. Interestingly, the VA treatment significantly decreased GAPDH expression in comparison to single treatments, suggesting differential drug efficacy between the two cell lines. HNRNP expression patterns revealed significant modulation by LS combination in both MOLM-13 and NOMO-1 compared to single treatments. While RU affected MOLM-13 in comparison to ruxolitinib and NOMO-1 in comparison to ulixertinib. The VA and VC treatments resulted in non-significant changes, indicating a more potent effect of LS and RU. RPL expression analysis showed that LS and RU treatments significantly modulated expression in MOLM-13 and NOMO-1. LS treatment led to a significant decrease in RPL expression (p = 8.13e-13), the same pattern observed in MOLM-13, RU treatment.

Across both zebrafish xenograft models, the LS and RU combinations consistently achieved the most significant modulation of gene expression in comparison to single treatments. In MOLM-13, LS and RU treatments resulted in substantial decrease of GAPDH, HNRNP, and RPL genes, indicating potent cellular activity suppression. The venetoclax-based combinations (VA, VC, and VR) displayed moderate effects, with VA demonstrating the strongest effect among them.

These results suggest that LS and RU combinations exert superior anti-leukemic effects, potentially through enhanced regulation of key metabolic and structural genes *in vivo*. The implications of these findings underscore the potential of these combinations in targeting specific mutations such as *FLT3, TP53, HLA-B* and *MUC4* associated with treatment resistance in AML, paving the way for improved therapeutic strategies in clinical settings. In conclusion, the differential gene expression patterns observed across treatments and cell lines highlight the complexity of drug responses in AML. The LS and RU combinations emerge as promising therapeutic strategies, demonstrating robust modulation of gene expression in both MOLM-13 and NOMO-1 xenografted embryos. These findings provide valuable insights into the molecular mechanisms underlying drug efficacy and resistance, informing the development of targeted therapies for AML.

## Discussion

The advent of targeted therapies, such as venetoclax, a selective BCL-2 inhibitor, has marked a paradigm shift in AML treatment, especially for patients who are older or unfit for intensive chemotherapy ^59,60^. Despite its clinical success, resistance to venetoclax-based therapies poses a substantial hurdle, often associated with poor prognostic mutations like *TP53* and *FLT3* ^35^. This highlights an urgent clinical need for novel therapeutic strategies that can effectively circumvent such resistance mechanisms, especially given the heterogeneity and complexity of AML’s genetic landscape.

In this study, we evaluated the efficacy of two novel combination therapies— LY3009120 plus Sapanisertib (LS) and Ruxolitinib plus Ulixertinib (RU)—across a diverse panel of AML cell lines, using a multilayered experimental framework that integrated *in vitro* viability assays, proteogenomic comparison, and *in vivo* zebrafish xenograft models. Our findings provide compelling evidence that both combinations not only outperform or match the efficacy of existing venetoclax-based regimens but also exhibit distinct advantages in overcoming drug resistance, particularly in genetically defined high-risk AML subtypes. The superior efficacy of the LS combination against AML cell lines, particularly MOLM-16 and SKM, underscores the potential of targeting both the RAS-RAF-MEK-ERK (MAPK) and mTOR pathways in overcoming monotherapies in oncology ^61,62^. The robust decline in cell viability observed with the LS combination suggests a synergistic effect, likely due to the simultaneous disruption of multiple signaling cascades that are critical for leukemic cell proliferation and survival. Similarly, the RU combination, by targeting the JAK/STAT and ERK pathways, achieved comparable or superior efficacy relative to venetoclax-based therapies. *TP53* and *FLT3* mutations were recurrently observed in cell lines that responded poorly to venetoclax-based regimens but exhibited heightened sensitivity to LS and RU. Notably, MOLM-16 and NOMO-1—harboring *TP53, MUC4*, and *HLA-B* mutations—emerged as especially responsive to LS and RU, highlighting the promise of these combinations in overcoming venetoclax resistance linked to adverse genetic profiles.

The strong response of MOLM-13 (*FLT3*-mutant) to LS also underscores the therapeutic potential of this regimen in *FLT3*-driven AML subtypes. These observations suggest a synergy between drug mechanism and mutational landscape, and support the stratification of AML patients based on actionable mutations to guide therapy selection. Interestingly, our findings also resonate with emerging perspectives that move beyond traditional mutation-centric models. A recent study by Jayavelu et al. proposed a reclassification of AML subtypes based on proteomic expression profiles rather than strictly genomic features and emphasized the role of mitochondrial metabolism in AML progression and drug resistance ^52^. Their work identified Mito-high subtypes as being associated with poorer outcomes and therapy resistance. While our dataset is not large enough to independently confirm the Mito-high/Mito-low hypothesis, our data compellingly align with it: Mito-high AML cell lines exhibited enhanced sensitivity to LS and RU. This suggests that metabolic profiling, alongside genomic analysis, could serve as a valuable biomarker for predicting treatment response. In particular, LS displayed exceptional efficacy in Mito-high, *FLT3*-mutated MOLM-13 cells, suggesting that dual inhibition of RAF and mTOR may be particularly well-suited to metabolically active AML subtypes.

Notably, these combinations were not rationally designed based on genetic alterations alone. Instead, they were derived through data mining of phenotypic drug response patterns, illustrating the utility of leveraging nominal-level screening to uncover hidden therapeutic synergies ^30,63,64-65^. Interestingly, we observed a correlation between mutational backgrounds and cell viability outcomes, which reinforces the validity of our approach.

The zebrafish xenograft model provided key translational validation for our *in vitro* findings. Both LS and RU significantly reduced leukemia burden in vivo, as evidenced by decreased fluorescence signals in MOLM-13 and NOMO-1 xenografted embryos. Importantly, these therapeutic effects were achieved with minimal developmental toxicity, as reflected by modest and statistically non-significant reductions in embryo length. This favorable efficacy-to-toxicity ratio supports the continued development of LS and RU regimens as viable treatment options. Furthermore, gene expression analyses in xenografted embryos reinforced the specificity and impact of these treatments. Two-way ANOVA revealed significant treatment-dependent gene expression modulation, particularly under LS and RU treatment, emphasizing the context-specific therapeutic effects of these combinations in distinct AML subtypes.

Our findings advocate for a shift toward rationally designed, genotype-guided combination therapies in AML. The LS and RU regimens not only exhibit strong cytotoxicity *in vitro* and *in vivo*, but also demonstrate efficacy in overcoming key resistance mechanisms associated with poor-prognosis mutations such as *TP53* and *FLT3*. By integrating multiomic analyses with functional drug screening, we demonstrate a blueprint for precision oncology in AML.

Overall, incorporating diverse biological data across different phenotypical and molecular levels—such as drug response, protein expression profiles, and genomic data— enables a more comprehensive identification of the unique vulnerabilities of individual tumors ^65–68^. Additionally, clinical trials incorporating mitochondrial profiling and genetic stratification could accelerate the translation of LS and RU into effective, personalized AML therapies. This integrative approach holds significant potential for improving treatment outcomes, particularly for patients with complex mutational landscapes. By uncovering actionable targets that may not be apparent through genomic analysis alone, this strategy paves the way for the development of more effective and personalized treatment protocols.

## Method

### Cell culture

Human myeloid leukemia cell lines (MOLM-16, SKM-1, MV4-11, MOLM-13, NOMO-1, KG-1, Kasumi-1, HL-60, PL-21, and ML-2) were obtained from (Leibniz Institute DSMZ-German Collection of Microorganisms and Cell Cultures GmbH). All cells were grown in RPMI-1640 (Gibco/Thermo Fisher Scientific, Waltham, MA, USA) supplemented with 10% or 20% FBS (Gibco), 1% L-Glutamine (Gibco), and 1% Penicillin-Streptomycin (Gibco). To prepare the cell suspensions for injection, cells were washed three times with PBS at 300 g for 3 minutes to remove any residual culture medium.

### Drug combination testing

The compounds were dissolved in dimethyl sulfoxide (DMSO) and dispensed on 384-well plates (Corning, Corning, NY, USA) for RTG and 24-well plates for zebrafish treatment. Drug concentrations were selected based on the previous study and cytotoxicity test for the zebrafish larvae. DMSO was used as a negative control by the concentration of 0.01% in RTG and 0.1% in the zebrafish experiment. *The drug concentrations are as below:* LY3009120 500nM, Sapanisertib 500nM, Ruxolitinib 3uM, Ulixertinib 3uM, venetoclax 50nM, azacitidine 300nM, cytarabine 1000nM

### Cell viability analysis using RealTime-Glo™

AML cell lines were seeded on 384-well plates (Corning) containing the desired concentration of drugs with three replicates. The number of cells per well was desired based on a linearity test as described in the manual for continuous-read experiment. In brief, cells were plated at increasing cell densities in the presence of 1X RealTime-Glo™ reagent, and cell viability was monitored by measuring luminescence every 12 hours for 72 hours. Based on the results the number of cells that remained linear in 72 h was selected (KG-1, MOLM-16, HNT, Kasumi-1 with 5000 cells per well and HL60, ML-2, MOLM-13, MN4-11, NOMO-1, PL-21, SKM with 1500 cell per well). Following the linearity test, the desired number of cells per well was adjusted to a final volume of 25 ul and were incubated with compounds for 72h at 37°C and 5% CO2. Cell viability was assessed using the CellTiter-Glo assay (Promega), and the luminescence signal was measured using a PHERAstar FS plate reader (BMG LABTECH, Ortenberg, Germany). Cell viability was measured at timepoints of 0, 12, 24, 36, 48, 60, and 72h.

### Zebrafish larvae microinjection and drug administration

For imaging purposes, cells were stained using CellTrace Far Red (Invitrogen, Carlsbad, CA, USA) according to the manufacturer’s instructions. At the final step of dying cells, cells were then suspended in media at a concentration of 50 × 10^3^ cells/μl. For microinjection, 4 nL of this suspension was injected, resulting in approximately 200 cells per embryo. The age of embryos is indicated as hours post-fertilization (hpf) and days post-fertilization (dpf). The larvae were grown at 28.5 °C in an embryonic medium (5 mM NaCl, 0.17 mM KCl, 0.33 mM CaCl2, and 0.33 mM MgSO4; Sigma-Aldrich). Two dpf, wild-type zebrafish (Danio rerio) larvae (from AB strain) were used. Embryos were dechorionated, anesthetized with 0.04 % tricaine, and microinjected through the yolk sac with 4 nL of pre labelled cells. 11200 cells were injected per fish. Fish were transferred into a 24-well plate (five fish per well) containing 1.5 mL of fresh embryonic medium with desired drug concentrations, and incubated for three days at 34 °C. The zebrafish larvae were treated with single and combinations as bellow concentration: LY3009120 500nM, Sapanisertib 500nM, Ruxolitinib 3uM, Ulixertinib 3uM, venetoclax 50nM, azacitidine 300nM, cytarabine 1000nM. Drug concentrations were chosen based on our previous in vitro drug testing as well as on the cytotoxicity test for the zebrafish larvae. Drugs were diluted in the embryonic medium and DMSO was used as the negative control. Twenty fish were used for each drug, where nine fish were pooled together to provide a sufficient signal during PCR amplification and the rest were used for imaging. Fish microin-jection and experiments were conducted at the Zebrafish Unit (University of Helsinki) and approved by the ethical permission from the regional state administrative agency (ESAVI/13139/04.10.05/2017).

### Zebrafish Fixation and imaging

Following 72h treatment, larvae were collected and fixed with 4 % paraformaldehyde and mounted on glass slides in SlowFade™ Gold antifade reagent (Invitrogen) for imaging. Zebrafish were imaged with a Zeiss Axio Imager (Carl Zeiss AG, Oberkochen, Germany), with 20x lens and zstack interval 12,5 μm and tiles. Then the tumor intensity was analyzed using ImageJ. Imaging was performed at the Biomedicum Imaging Unit, the University of Helsinki, supported by the Helsinki Institute of Life Science (HiLIFE) and Biocenter Finland. All experiments were repeated three times independently, with 20 larvae per group.

### Primer design and real-time quantitative PCR

Sequences of candidate genes were searched from GenBank for PCR primer designing using NCBI with Tm 59–62°C, primer length 18–22 bp, guanine-cytosine content 50–60%, and amplicon length 80–220 bp. The secondary structures of primers were checked using Gene Runner. We designed three primers for human genes and four for zebrafish (Supplementary Table S1). Quantitative PCR was performed using Lightcycler-FastStart DNA Master SYBR Green 1 and Lightcycler480 (Roche) according to the manufacturer’s instructions. The qPCR was performed in 10 μl with a primer concentration of 1 μM, 10 ng cDNA, and 1× SYBR Green qPCR Master Mix (Roche). The amplification program consists of an initial denaturation step at 95° C for 12 min and 40 cycles of denaturation at 95° C for 10s, annealing at 61°C for 15s and extension at 72° C for 15s. A total of two biological replicate samples were analyzed in duplicates (technical duplicates).

Since the PCR efficiency of the primer pair limitedly affects the analysis of gene expression stability across samples, the PCR efficiencies of the primers used were above 1·8, as determined by qPCR with serially diluted pooled cDNA samples (10^6^, 10^5^, 10^4^, 10^3^ dilutions), which was slightly beyond the recommended range 1,90 to 2,05.

### Statistical analysis

Values are given as means ± standard deviations. To determine the statistical significance, we performed non-paired Student’s T-test, and one-way analysis of variance (ANOVA) followed by Tukey’s post-hoc test. We set statistical significance to P ≤ 0.05. The mutation lists for cell lines are downloaded from DepMap.

## Supplementary data

**Supplementary table S1.** Primer sequences used for qPCR analysis.

**Supplementary table S2.** qPCR analyzed using two-way Classical ANOVA.

**Supplementary table S3.** qPCR analyzed using ANOVA followed by post hoc pairwise comparisons.

**Supplementary Figure S1.** Detailed Cell viability results for all 10 Aml cells within all treatments from 12-72h. Cell viability (%) was monitored over 72 hours in 10 AMl cell lines (KG-1, MOLM-16, HNT, Kasumi-1, HL-60, ML-2, MOLM-13, MN4-11, NOMO-1, PL-21, SKM-1). Error bars indicate standard deviations. Abbreviations: LY3009120 (L), Sapanisertib (S), Ruxolitinib (R), Ulixertinib (U), Venetoclax (V), Azacitidine (A), Cytarabine (C), LY3009120 & Sapanisertib (LS), Ruxolitinib & Ulixertinib (RU), Venetoclax & Azacitidine (VA), Venetoclax & Cytarabine (VC), Venetoclax & Ruxolitinib (VR).

## Acknowledgments

We extend our gratitude to the Biomedicum Imaging Unit and the Zebrafish Unit at the University of Helsinki, Finland, for their invaluable support throughout this study. We are particularly thankful to Mr. Henri Koivula for his expert assistance with zebrafish larvae microinjections. Additionally, we sincerely appreciate Dr. Sadegh Azimzadeh for his valuable advice on data analysis. This study was financially supported by the Research Council of Finland [Grant 332454 to M.J.] and the Jane and Aatos Erkko Foundation [Grant 220031 to M.J.].

