## Supplementary figures and images for "Targeting AML Resistance with Two Novel Combinations Demonstrate Superior Efficacy in *TP53, HLA-B, MUC4* and *FLT3* mutations"

### Supplementary Fig 1

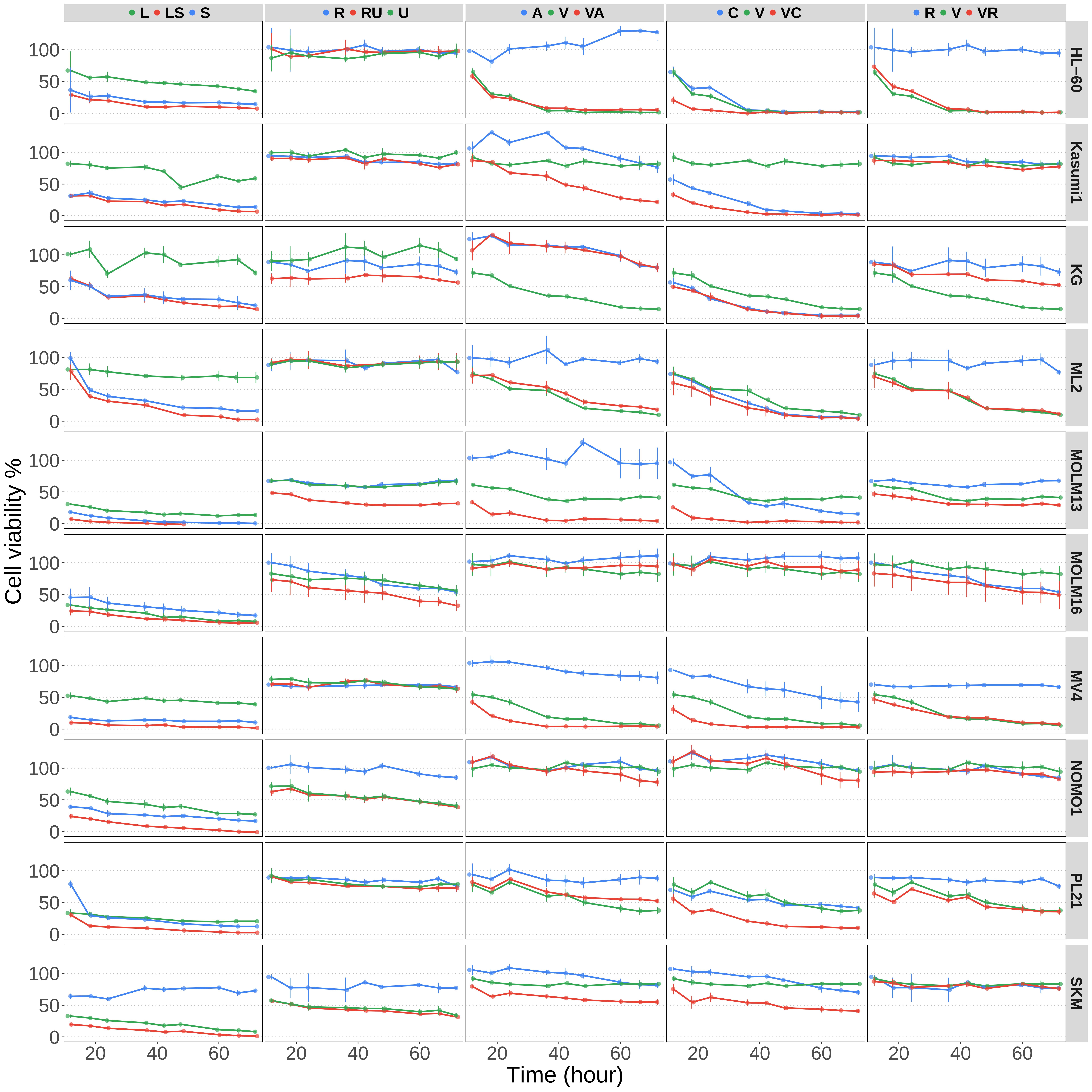
